# Multi-site phenotypic characterisation of Kisumu colonies for comparison of insecticide susceptibility between testing centres

**DOI:** 10.64898/2026.07.21.739746

**Authors:** Gemma Francesca Harvey, Giorgio Praulins, Sylvain Akoupo, Godwin Kwame Amlalo, Selina Anthony, Salum Azizi, Koama Bayili, Edmond Bernard, Vincent Corbel, Kounbobr Roch Dabire, Abdoulaye Diabate, Stéphane Duchon, Joannitta Joannides, William N. Kisinza, Benjamin G. Koudou, Aneth Mahande, Alphaxard Manjurano, Johnson Matowo, Benson Mawa, Sarah J Moore, Ahmadi B Mpelepele, Gift Mwaanga, Victor Mwingira, Boris N’dombidje, Raphael N’Guessan, Corine Ngufor, Eric Ochomo, Wema S. Saidi, Kochelani Saili, Limonty Simubali, Jennifer C Stevenson, Magellan Tchouakui, Yvan Fotso Toguem, Tyler Walker, Jessica Williams, Rosine Z. Wolie, Charles Wondji, Nick Yalla, Akua Obenewaa Danquah Yirenkyi, Julien Z. B. Zahouli, Frank Mechan, Alexandra Wright, Rosemary Susan Lees

## Abstract

Testing of insecticides and insecticide-based vector control tools relies on the use of bioassays whereby insects are exposed and mortality, or another relevant entomological endpoint, is measured. To be able to compare results between products and across time and space standard laboratory strains are used. Insecticide-susceptible strains are often used as comparators in studies involving insecticide-resistant populations. ‘Kisumu’ is a long-established, widely-used insecticide-susceptible strain of *Anopheles gambiae*. In this study, 16 facilities across different countries exposed their Kisumu colonies aged 3-5 days to four insecticides (permethrin, alpha-cypermethrin, DDT and pirimiphos-methyl) at a range of concentrations in WHO tube assays to assess susceptibility. Insecticide-treated filter papers were made at the Liverpool School of Tropical Medicine and sent to testing centres, along with a standard protocol and testing conditions requirements, to standardise testing as much as possible. A first observation was that DDT results were affected by the use of Risella oil, the WHO-recommended carrier oil but immiscible in acetone, leading to uneven coating of papers and increased variability in bioassay results (40x variance between colonies). Bioassay data generated by the testing facilities suggest a wide range of susceptibility to different insecticides between colonies of Kisumu: LC_50_ values between colonies differed, though not significantly, by up to 4x for permethrin, 6x for pirimiphos-methyl, and 15x for alpha-cypermethrin. Some colonies are classified as insecticide resistant based on the WHO definition of <98% mortality on exposure to a discriminating concentration. The assumption that Kisumu is a standard susceptible strain, consistent between facilities is challenged, which has implications when comparing results between studies. We propose genotypic analysis of the colonies to identify signs of resistance markers indicative of contamination from resistant strains or wild mosquitoes, and generation of guidance to support the maintenance and monitoring of susceptible mosquito strains.

## Introduction

During the development, evaluation, and monitoring of vector control tools, particularly those containing insecticides, bioassays are used to measure their efficacy against the target species. In these assays, vectors are exposed to the insecticide or tool and outcomes such as mortality, or other relevant entomological endpoint, is measured to determine the tool’s effectiveness. For example, to demonstrate that an insecticide treated net (ITN) designed to reduce malaria transmission, is effective, a WHO cone test ^1^ may be used to test efficacy against *Anopheles gambiae* mosquitoes. Because bioassays rely on the use of a living test system, they are subject to many sources of variability or ‘noise’ which may bias results or affect reproducibility ^2–4^. To minimise the variability and maximise the ability to compare results between experiments performed at different facilities or ensure consistency of results over time the inputs into the assay need to be considered and standardised where possible, or characterised where this is not possible to help with interpretation of the results.

One of the key inputs to a bioassay is the mosquito test system for testing insecticides or vector control tools, and the genetic background and phenotypic response of an insect population will affect the results obtained. Characterising the mosquito strain used is valuable in being able to interpret results ^5^, and standard laboratory strains are often used to give standardisation over time ^6^ and between facilities. Insecticide-susceptible strains are often used to collect baseline data in studies, for example when screening for new insecticidal compounds ^7^ or developing ^8^ or evaluating ^5^ new products and used as comparators in studies involving insecticide-resistant mosquito populations ^9,10^. Kisumu is one such widely-used reference strain. According to BEI Resources, who hold the KISUMU1 colony and supply eggs or frozen material on request, *Anopheles gambiae* KISUMU1 stock was established by G. Davidson in 1975 in Kisumu, Kenya, Africa, and selected by French research institutes LIN and ORSTOM for permethrin susceptibility. The colony was contributed to CDC for the BEI Resources collection (formerly MR4) by Vincent Corbel, Institut de Recherche pour le Développement (IRD) in 1998.

Colonies of the Kisumu strain are maintained at many testing facilities, but the origin and exact nature of the strain held by these facilities may be different. Although some may have been established from material obtained from BEI Resources directly, others will have been established from eggs supplied from another facility or may have been colonised independently from the Kisumu region. Colonies all thought to be the same Kisumu strain may, in fact, differ considerably between facilities and even over time within the same facility for several reasons other than origin. Colonies may vary in their general fitness and insecticide susceptibility as a result of selection pressures applied by rearing methods, genetic bottlenecks when colonies were established, genetic drift over time, contamination with material from other strains which may include insecticide resistance alleles, or inadvertent selection pressure applied as a result of insecticide contamination.

Evidence from one example, the 2017–2021 WHO multi-centre study to establish discriminating concentrations (DC) for a range of insecticides ^11^ and subsequent work, shows that this assumption is not always supported. Multi-centre studies to establish discriminating concentrations for insecticide resistance monitoring have traditionally relied on well characterised, fully susceptible mosquito strains, assuming these provide a stable and comparable biological baseline across laboratories. Where possible, recognised reference strains such as Kisumu for *An. gambiae* and FANG for *An. funestus* have been used, while for other species, including *Ae. aegypti* and *An. stephensi*, different susceptible strains have been used according to local availability ^12,13^. This approach assumes that susceptible strains of the same species, and particularly those sharing the same strain name, are biologically equivalent between centres.

During the 2017-2021 study, Bayesian binomial analyses with flexible concentration-response functions demonstrated substantial inter-laboratory variability in lethal concentration estimates for the same insecticide species combinations. Within bioassay variability was generally low, indicating that the WHO bottle bioassay is robust and that observed differences arise primarily from biological variation between laboratory colonies rather than methodological artefacts. Variability was greatest at the upper end of concentration-response curves, with LC_99_ estimates showing markedly higher uncertainty than lower effect levels ^13^.

In the WHO-sponsored 2024–25 multi-centre study to establish DC for broflanilide and isocycloseram ^12^, pronounced inter-site variability was also observed, even when testing nominally identical susceptible strains, particularly Kisumu. Independent testing by the U.S. President’s Malaria Initiative (PMI) Evolve Project and Abt Global, using the same bottle bioassay methodology, assessed the susceptibility of colonised and field-sampled *An. gambiae* s.s. and *An. gambiae* s.l. to broflanilide across several sub-Saharan African countries. This work tested 3, 6, 9 and 18 µg/bottle; almost complete mortality was observed at 18 µg/bottle, with lower mortality in some tests at the lower concentrations, supporting the proposed preliminary discriminating concentration of 18 µg/bottle for *An. gambiae* ^14^.

Independent bottle bioassay testing generated a DC of 11.91 µg per bottle ^8^, consistent with another study using Kisumu with 500 ppm MERO ^15^. The WHO multi-centre laboratory study ultimately proposed discriminating concentrations of 10 µg/bottle for *An. gambiae*, 15 µg/bottle for *An. funestus*, and 25 µg/bottle for *An. stephensi*, broadly aligning with the 18 µg/bottle now proposed by Mitsui ^14^. Taken together, these findings show that susceptible laboratory strains cannot be assumed to be directly comparable across centres. Even colonies carrying the same strain designation can exhibit materially different concentration- response behaviour, with direct consequences for discriminating concentration selection.

Many facilities, particularly those with GLP certification, will perform routine profiling or characterisation of their reference colonies: for example, the LITE facility at the Liverpool School of Tropical Medicine has published data and protocols describing their genetic and phenotypic profiling of colonies including Kisumu, along with details of the origin of their colonies ^6^. However, a consultation with manufacturers of insecticide-based vector control tools ^16^ highlighted that the maintenance of an insecticide susceptible colony still represented a challenge in the generation of robust data when performing laboratory testing of vector control products. It is rare that data describing the susceptibility profile or other characteristics of the susceptible mosquito strains used for testing are reported in publications, as is recommended for insecticide resistant colonies used for, as an example, ITN bioefficacy testing ^24^. One of the recommendations emerging from the WHO multi-centre study on determination of insecticide discriminating concentrations for monitoring of resistance in Culex ^17^, was that participating laboratories in future similar studies should characterise the test strains using, for example, synergist bioassays and molecular assays, to improve quality control of the mosquito colonies. This emerged from an investigation into the reason behind elevated tolerance to pyrethroids in a ‘susceptible’ reference strain held at one testing centre, which identified the presence of a kdr mutation which was conferring resistance to pyrethroids. Had the strains been characterised before the study, this issue would have been identified and the strain excluded from the study, saving wasted testing resource.Similarly, whilst many scientific publications each year present data generated using the ‘Kisumu’ strain, the actual mosquitoes used for testing in each study or at each facility may be very different in fitness and susceptibility status and, therefore, results generated may not be comparable.

The aim of this study was to characterise Kisumu colonies maintained in as many facilities across the globe as possible by assessing their susceptibility to four representative insecticides: permethrin, alpha-cypermethrin, DDT and pirimiphos-methyl. Filter papers were treated with the discriminating concentrations (DC) of each insecticide ^18^, at which mortality in a susceptible population would be expected to be >98%, and a serial dilution expected to produce a dose response for each colony which could be compared. Sixteen test facilities used these papers in WHO tube bioassays with their Kisumu colony, with the goal of determining how comparable results were to test the assumption that all Kisumu colonies are equivalent and whether using this standard colony for bioassays truly removes the mosquito test system as a source of variability in testing.

## Methods

### Identification of participating laboratories

All the laboratories known to the authors who may hold a colony of Kisumu and with the ability to perform standard bioassays were contacted to invite them to participate in the study, on confirmation that they held a Kisumu colony. Using a snowball approach, we identified additional laboratories to invite.

### Treatment of insecticide papers

Preliminary experiments were performed using the *Anopheles gambiae* Kisumu strain, originally established from Kisumu, Kenya, and has been at the Liverpool School of Tropical Medicine (LSTM) since 1975. The Liverpool Insect Testing Establishment (LITE) routinely profiles the colony against insecticides of different classes to confirm its susceptibility ^6^.

Concentration–response curves were generated using WHO tube assays with filter papers treated with 1x, 1/2x, 1/4x, 1/8x, 1/16x, and 1/32x of the discriminating concentration (DC) of permethrin, pirimiphos-methyl, dichlorodiphenyltrichloroethane (DDT), and alpha-cypermethrin. These papers were tested in a single replicate, and the results used to further refine the concentration ranges. Upper and lower limits were adjusted where necessary, and intermediate concentrations were modified to maximise coverage of the full response curve while maintaining clear separation between adjacent concentrations. The final concentrations (Table 1) were selected to allow for potential differences in colony responses between centres to be detected.

**Table 1.**
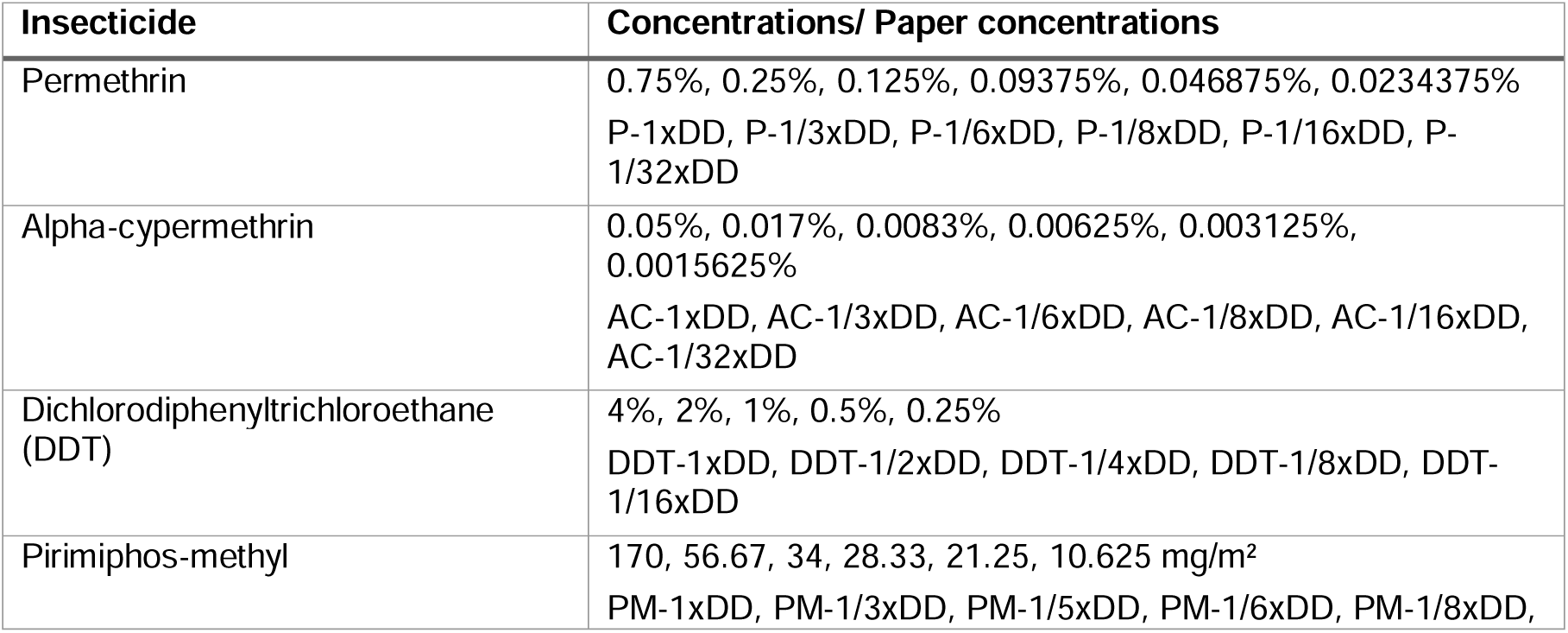
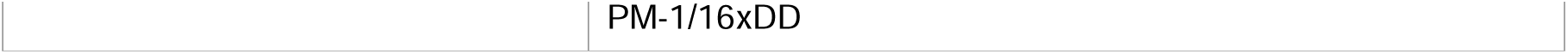
Concentrations tested for each insecticide (one paper for each concentration).

Test papers were made up according to the WHO SOP ^19^. Pyrethroid papers were treated with silicone oil, DDT papers treated with Risella oil, and pirimiphos-methyl papers made without oil. For each insecticide, one test paper was made up per concentration per testing site. Sixteen sites were sent insecticide-treated papers, along with control papers: pyrethroid control (for permethrin and alpha-cypermethrin assays), acetone-only control (pirimiphos-methyl assay) and organochlorine control (DDT assay).

### Phenotypic characterisation of Kisumu colonies: determining their true susceptibility status

The 16 testing sites were asked to follow the WHO tube bioassay SOP ^20^, exposing 25 3-5 day old non-blood fed adult females to treated papers for 60 minutes in environmentally controlled testing rooms (27 +/- 2°C and 75% +/- 10% relative humidity). Sites were advised to use 50 mosquitoes per control (2 tubes) and 100 mosquitoes per test (4 tubes) on each treated paper. They were asked to record knockdown per concentration at 15, 30, 45 and 60 minutes and measurements of 24-hour mortality were also recorded.

### Data analysis

Concentration–response curves and LC values were estimated using a Bayesian five-parameter logistic framework of Kont et al. ^21^ implemented in Stan via the rstan interface (R version 4.4.3) and with the same framework as applied in WHO multi-centre studies ^13^. The number of mosquitoes responding out of the total tested at each concentration was treated as binomially distributed, weighting each observation by the number assayed and accommodating variation in mosquito numbers between bioassays. Concentration was square root transformed prior to fitting, and background mortality was estimated within the model rather than by post-hoc Abbott’s correction. A separate curve was fitted to each site for each insecticide, with four chains of 10,000 iterations; convergence was confirmed by the Gelman–Rubin statistic (R < 1.01) together with zero divergent transitions. Prior distributions and the fixing of the lower and upper asymptotes follow Kont et al. ²¹. LC values are reported as posterior medians with 95% credible intervals, taken as the 2.5^th^ and 97.5^th^ percentiles of the posterior.

Between-site LC_50_ fold differences were calculated for each insecticide as the largest site LC divided by the smallest, expressing each site’s LC relative to the most susceptible site. If mortality was observed at the discriminating concentration (the highest concentration tested), the average proportional 24-hr mortality was assessed against a <98% threshold indicative of resistance ^18^.

Prior to curve fitting, replicates with control mortality of 20% or higher were excluded, and any data point recording greater than 100% mortality was removed. One further replicate (replicate 1, site M, pirimiphos-methyl) was excluded because mortality was 0% at the lowest doses and 100% at the highest, inconsistent with replicates 2 and 3 and not contributing to a coherent dose–response.

Data were considered unsuitable for concentration–response analysis, and the corresponding site excluded for that insecticide, where either (i) fewer than four distinct concentrations were tested, too few to identify the five parameters of the logistic model, or (ii) mortality at the lowest concentration tested was ≥98%, indicating the assay did not bracket the LC and leaving no estimable rising portion of the curve. For these sites, scatter plots were produced to show the variability in mortality. Curves meeting these criteria were retained even where individual estimates were imprecise; such cases are reported with their full credible intervals.

To assess whether the relative susceptibility of the Kisumu strain to different insecticides was consistent across testing sites, we examined the rank correlation of site-level LC estimates between insecticides. For each insecticide, sites were ranked by their posterior median LC , and Spearman’s rank correlation coefficient (ρ) was calculated for every pair of insecticides. Correlations were computed pairwise, each comparison including all sites with an estimable LC for both insecticides in the pair (n = 13 for comparisons among DDT, alpha-cypermethrin and permethrin; n = 9 for comparisons involving pirimiphos-methyl, for which fewer sites yielded estimable LC values). A rank-based correlation was chosen because it compares the ordering of sites rather than absolute LC magnitudes, which differ by orders of magnitude between compounds, and because it is robust to the non-normal distribution of LC values. Two-sided p-values were obtained from the asymptotic t-approximation. To propagate LC uncertainty into the correlation, Spearman’s ρ was additionally recomputed across matched posterior draws of the site LC values, yielding a median ρ and 95% credible interval for each insecticide pair. No correction was applied since the analysis was hypothesis-generating rather than confirmatory, and the n value was small. Analyses were performed in R, and the correlation matrix was visualised as a heatmap using ggplot2.

To compare susceptibility between testing sites within each insecticide, the ratio of each pair of sites’ LC values was computed across matched posterior draws of the site-level estimates. Two sites were considered to differ where the 95% credible interval of this ratio excluded 1; sites whose credible intervals overlapped were regarded as statistically indistinguishable, providing a Bayesian equivalent of a pairwise significance test (visualised in the site LC panels, panel C of Figures 2–5).

**Figure 1.**
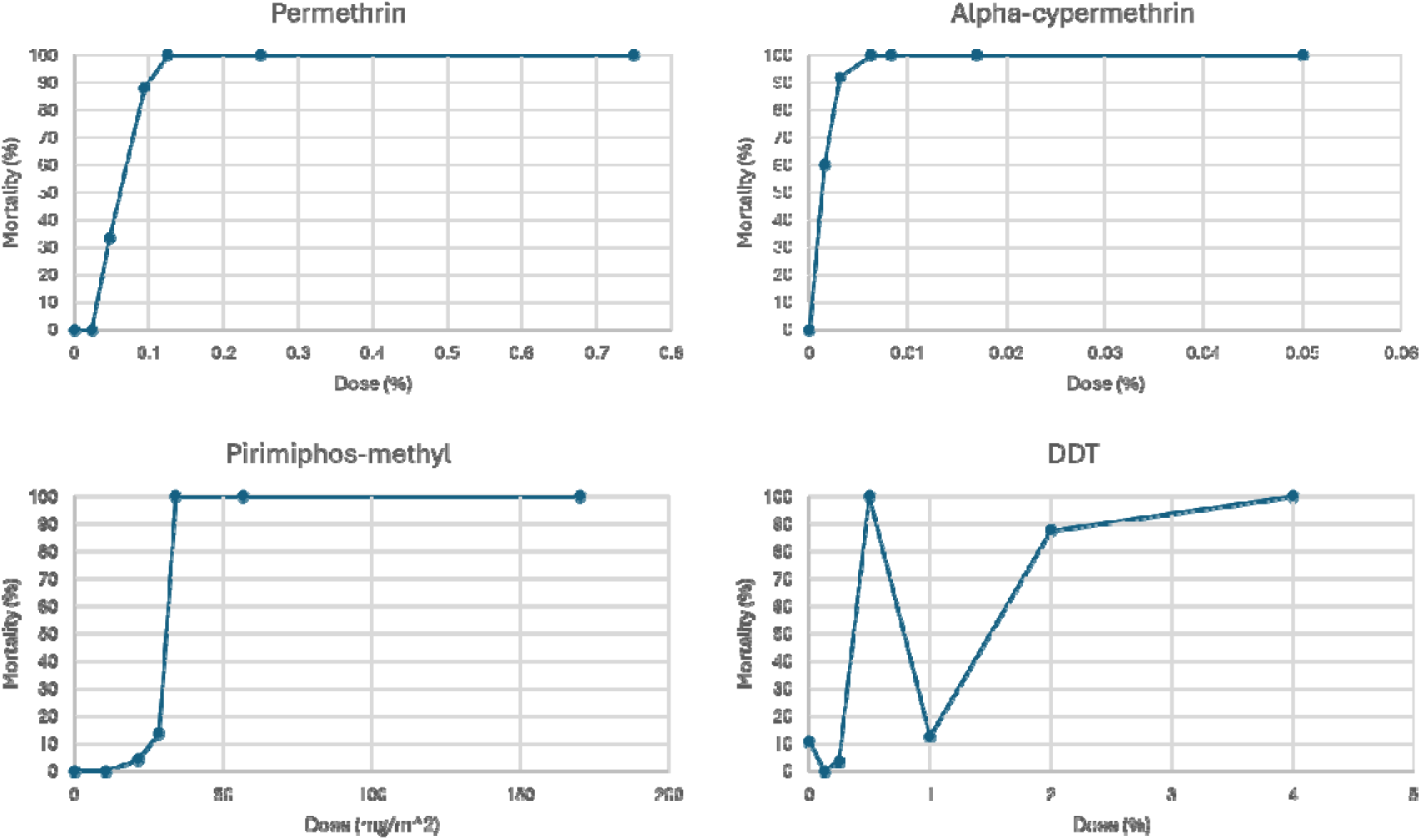
Results from a single replicate range finding experiment used to select concentrations for testing Kisumu colonies in participating testing facilities. The Kisumu colony held by the Liverpool Insect Testing Establishment (LITE) was exposed to filter papers treated with a range of concentrations on each insecticide in the WHO tube assay.

**Figure 2.**
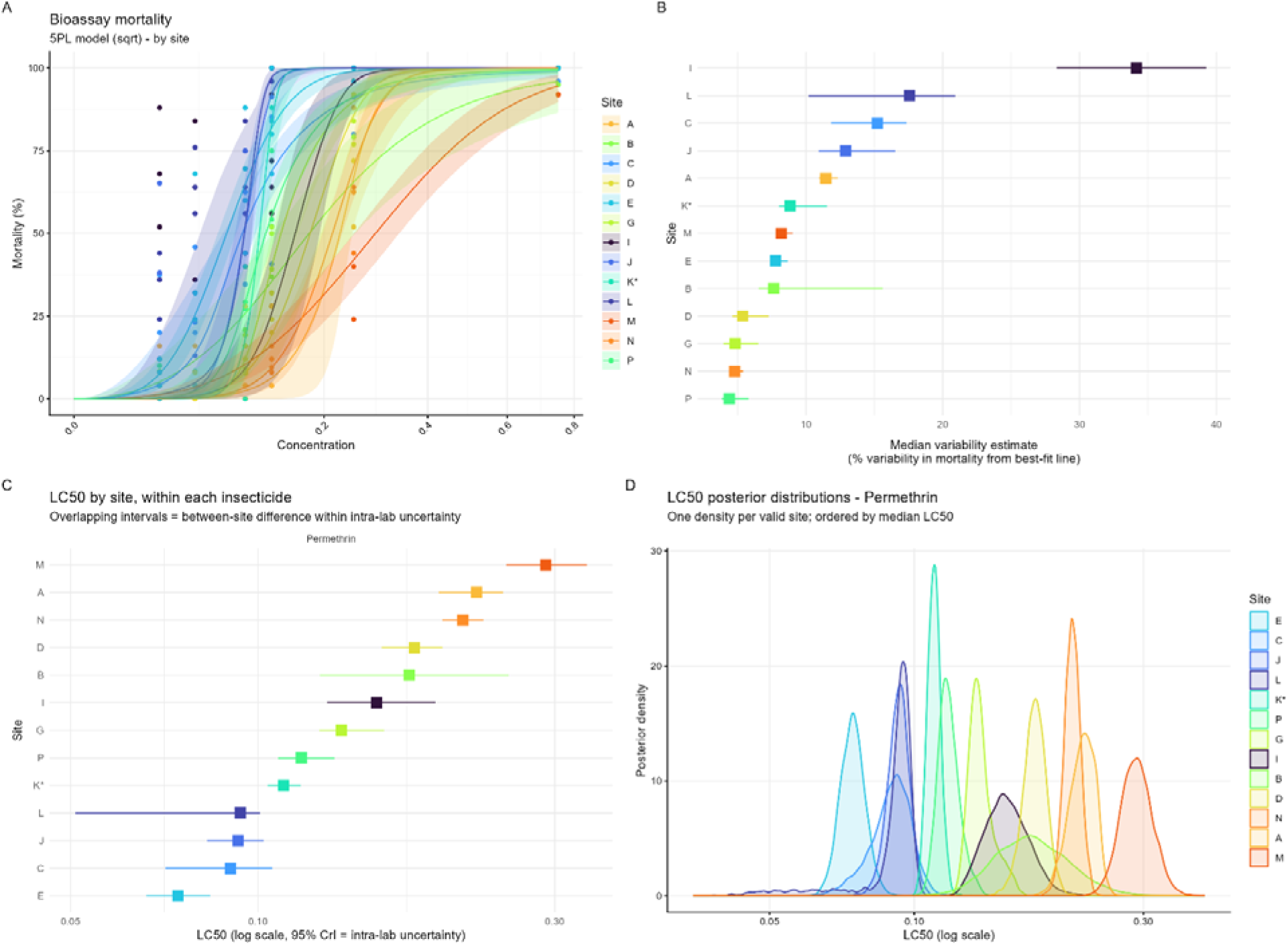
Inter-laboratory variability in permethrin susceptibility of Kisumu mosquitoes measured using the WHO tube bioassay. **(A)** Five-parameter logistic (5PL) concentration–response curves showing 24-hour mortality (%) across anonymised testing sites. Points represent observed mortality, lines represent fitted dose–response relationships, and shaded areas indicate 95% credible intervals. **(B)** Estimated unexplained mortality variability for each site. Squares indicate median estimates and horizontal lines show 95% credible intervals. **(C)** Site-specific LC_50_ estimates for permethrin, ordered by increasing LC_50_. Squares represent median posterior estimates, and horizontal lines indicate 95% credible intervals. **(D)** Posterior distributions of site-specific LC_50_ estimates, illustrating uncertainty associated with each site’s LC_50_ estimate.

**Figure 3.**
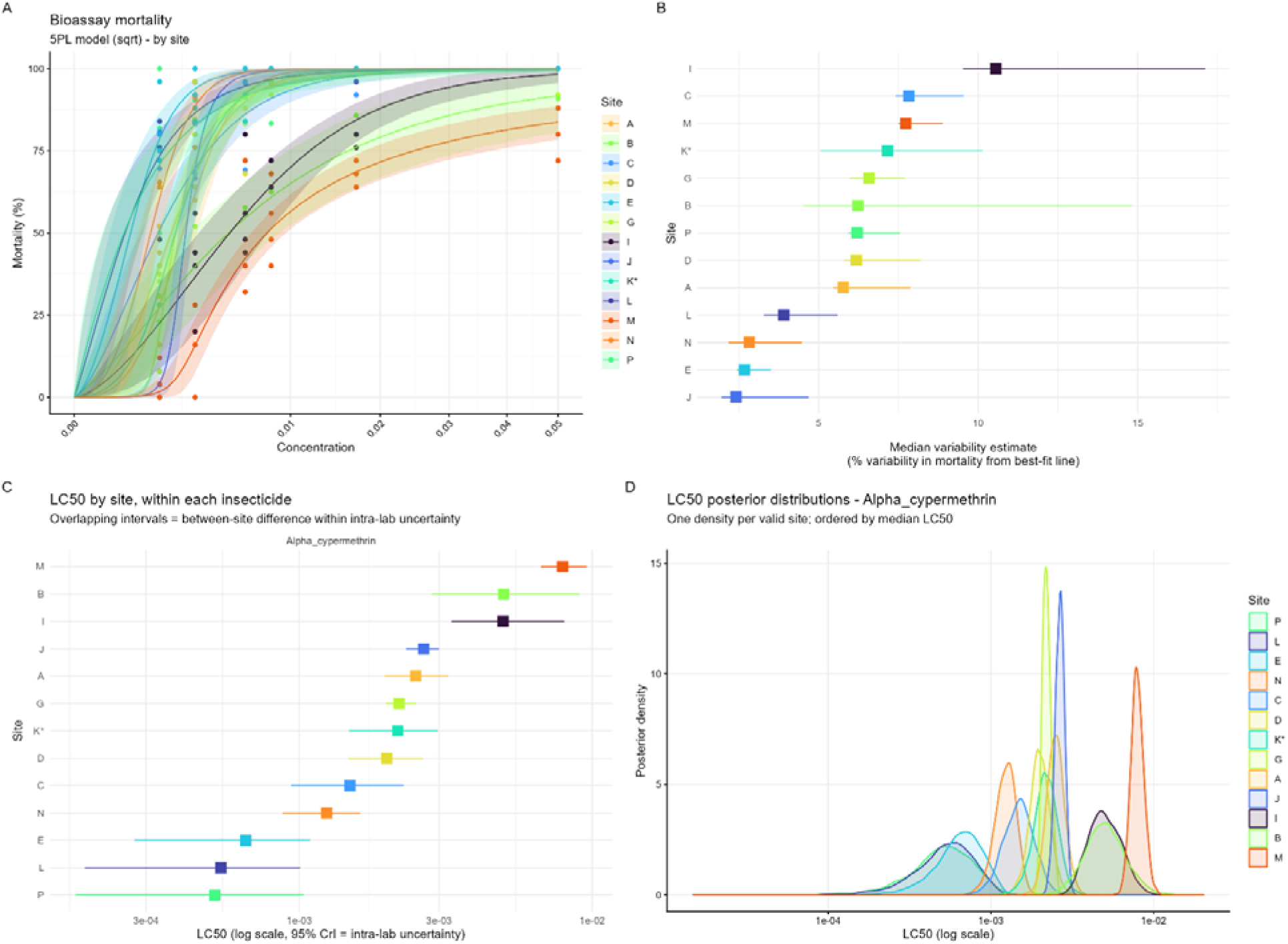
Inter-laboratory variability in alpha-cypermethrin susceptibility of Kisumu mosquitoes measured using the WHO tube bioassay. **(A)** Five-parameter logistic (5PL) concentration–response curves showing 24-hour mortality (%) across anonymised testing sites. Points represent observed mortality, lines represent fitted dose–response relationships, and shaded areas indicate 95% credible intervals. **(B)** Estimated unexplained mortality variability for each site. Squares indicate median estimates and horizontal lines show 95% credible intervals. **(C)** Site-specific LC_50_ estimates for alpha-cypermethrin, ordered by increasing LC_50_. Squares represent median posterior estimates, and horizontal lines indicate 95% credible intervals. **(D)** Posterior distributions of site-specific LC_50_ estimates, illustrating uncertainty associated with each site’s LC_50_ estimate.

**Figure 4.**
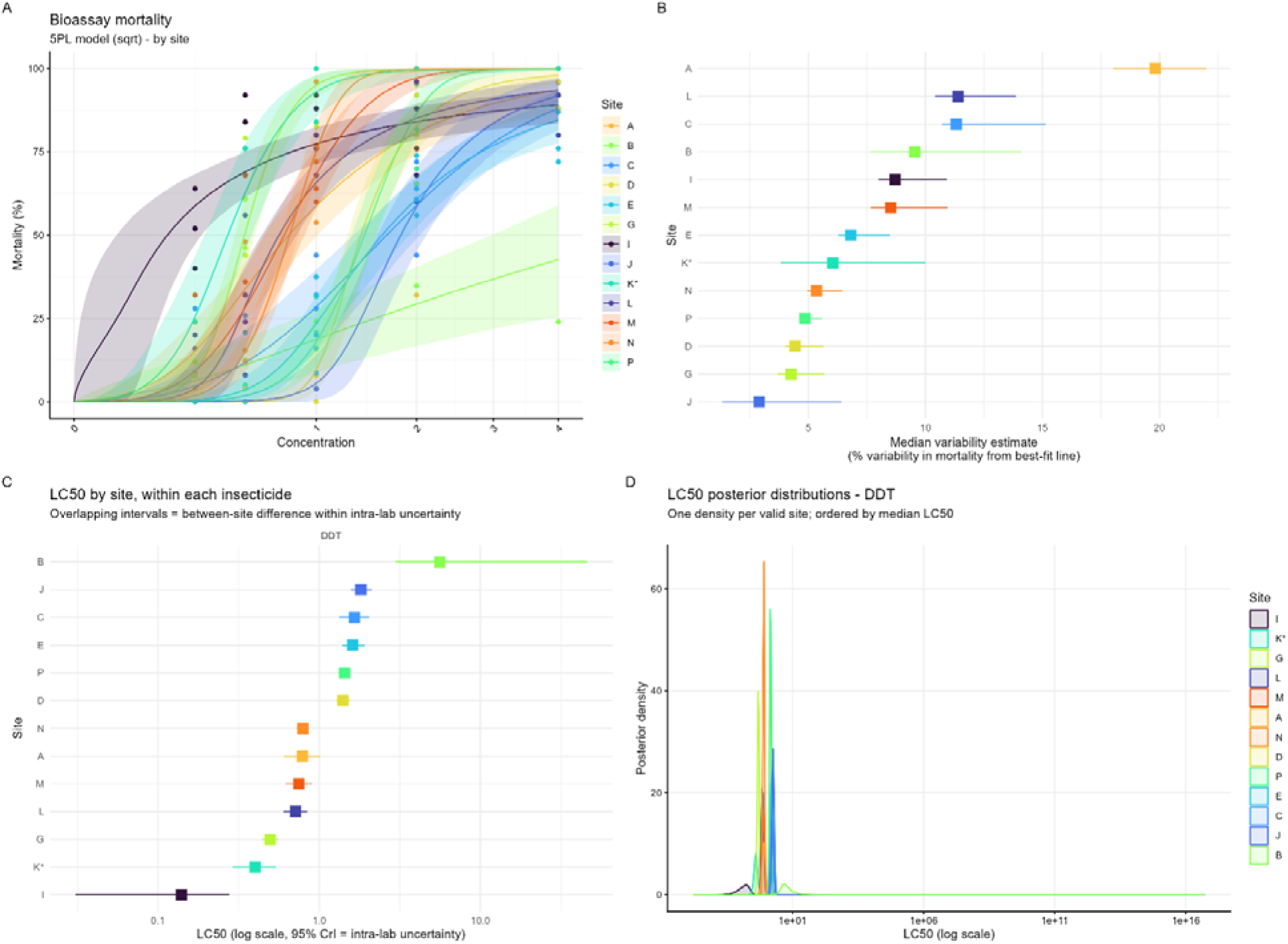
Inter-laboratory variability in Dichlorodiphenyltrichloroethane (DDT) susceptibility of Kisumu mosquitoes measured using the WHO tube bioassay. **(A)** Five-parameter logistic (5PL) concentration–response curves showing 24-hour mortality (%) across anonymised testing sites. Points represent observed mortality, lines represent fitted dose–response relationships, and shaded areas indicate 95% credible intervals. **(B)** Estimated unexplained mortality variability for each site. Squares indicate median estimates and horizontal lines show 95% credible intervals. **(C)** Site-specific LC_50_ estimates for DDT, ordered by increasing LC_50_. Squares represent median posterior estimates, and horizontal lines indicate 95% credible intervals. **(D)** Posterior distributions of site-specific LC_50_ estimates, illustrating uncertainty associated with each site’s LC_50_ estimate.

**Figure 5.**
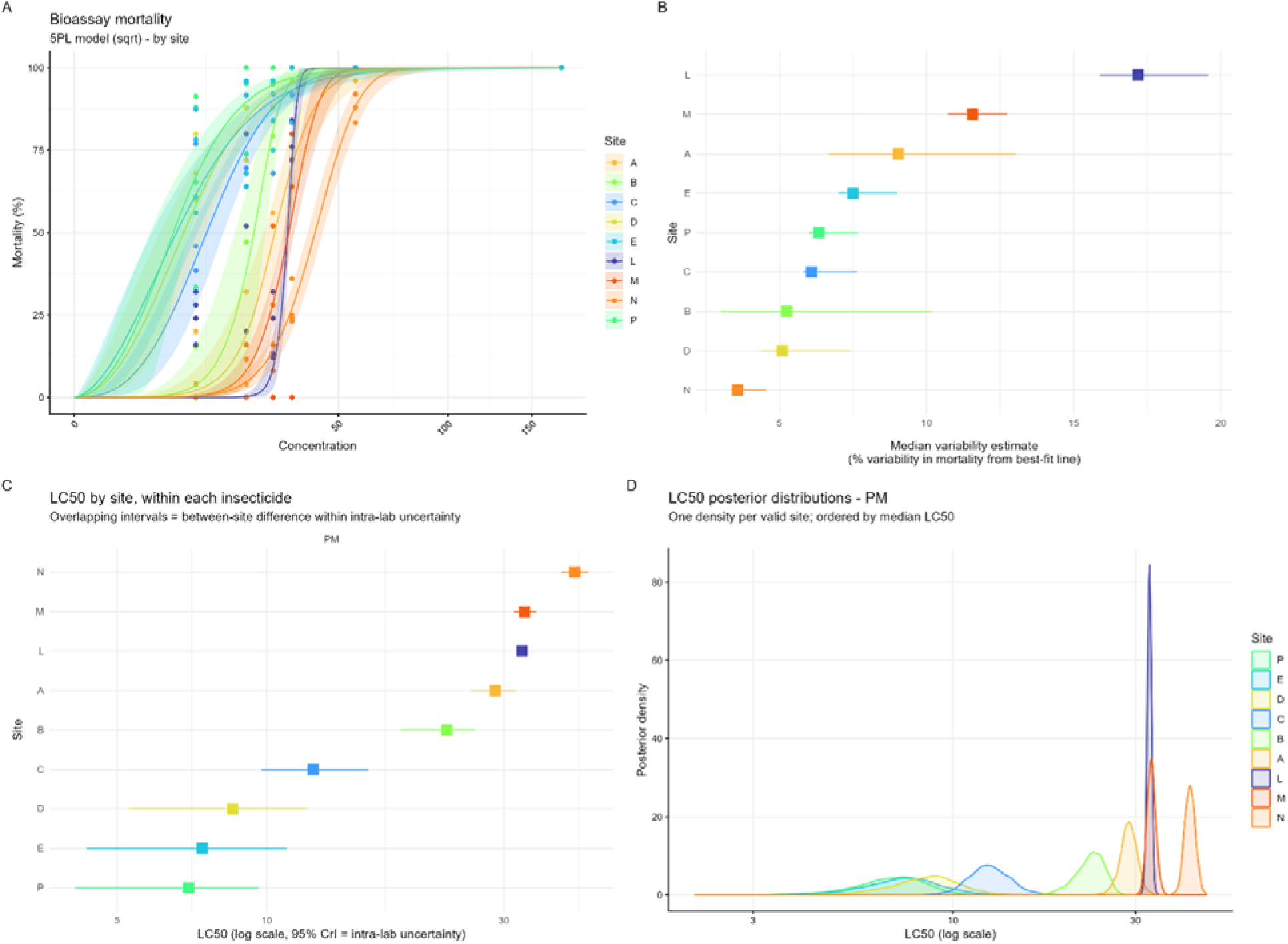
Inter-laboratory variability in Pirimiphos-methyl susceptibility of Kisumu mosquitoes measured using the WHO tube bioassay. **(A)** Five-parameter logistic (5PL) concentration–response curves showing 24-hour mortality (%) across anonymised testing sites. Points represent observed mortality, lines represent fitted dose–response relationships, and shaded areas indicate 95% credible intervals. **(B)** Estimated unexplained mortality variability for each site. Squares indicate median estimates and horizontal lines show 95% credible intervals. **(C)** Site-specific LC_50_ estimates for pirimiphos-methyl, ordered by increasing LC_50_. Squares represent median posterior estimates, and horizontal lines indicate 95% credible intervals. **(D)** Posterior distributions of site-specific LC_50_ estimates, illustrating uncertainty associated with each site’s LC_50_ estimate.

## Results

### Preliminary Concentration Range Testing

The initial concentration ranges tested confirmed that the selected concentrations were sufficient to span a full concentration-response range, producing mortality values from 0 to 100 percent. Minor adjustments were then made to the concentrations used for the study to improve separation between concentrations in terms of observed mortality. Lower concentrations were occasionally removed where multiple concentrations produced negligible mortality. For example, where both 1⁄16×DD and 1⁄32×DD resulted in less than 5 percent mortality, the 1⁄32×DD concentration was dropped. At the upper end of the range, some concentrations were shifted downwards to avoid testing multiple concentrations that consistently produced 100 percent mortality. For permethrin and alpha-cypermethrin, the 1⁄2×DD and 1⁄4×DD concentrations were therefore replaced with 1⁄3×DD and 1⁄6×DD, respectively.

For DDT, only the 1⁄32×DD concentration was removed. For pirimiphos-methyl, the entire concentration range was adjusted due to a sharp increase in mortality observed between the 1⁄4×DD and 1⁄8×DD concentrations. The revised range comprised 1⁄3×DD, 1⁄5×DD, 1⁄6×DD, 1⁄8×DD, and 1⁄16×DD, with the 1⁄32×DD concentration omitted. The DDT dose range showed an unexpectedly high mortality at the 1⁄8×DD concentration. During preparation of the papers, it was noted that Risella oil, used as the carrier for DDT, is immiscible in acetone and required repeated vortexing to ensure both the solvent and oil were transferred onto the filter paper. This made it more difficult to achieve even coverage compared with other insecticides. However, despite this the selected concentration range was considered sufficient to cover the full range of mortality.

### Participating laboratories and numbers of mosquitoes tested

A total of sixteen laboratories agreed to participate. These included product development laboratories run by manufacturers of vector control tools, contract research laboratories, and private and public research institutions.

A total of 8,569 Kisumu mosquitoes were exposed to permethrin, 8,559 to alpha-cypermethrin, 7,553 to DDT, and 8,309 to pirimiphos-methyl (Table 2), a grand total of 32,990 mosquitoes.

**Table 2.** Numbers of mosquitoes tested per test site, per insecticide. DDT = Dichlorodiphenyltrichloroethane, PM = Pirimiphos-methyl. For sites F, H and O an LC50 could not be calculated as only the control, and the lowest and highest concentrations were included in testing due to resource constraints

| Site | DDT | Permethrin | Alpha-cypermethrin | Pirimiphos-methyl | Subtotal per site |
| --- | --- | --- | --- | --- | --- |
| A | 500 | 400 | 400 | 400 | 1700 |
| B | 251 | 213 | 213 | 195 | 872 |
| C | 572 | 702 | 693 | 643 | 2610 |
| D | 450 | 525 | 525 | 525 | 2025 |
| E | 686 | 790 | 789 | 783 | 3048 |
| F | 290 | 319 | 316 | 311 | 1236 |
| G | 691 | 790 | 795 | 790 | 3066 |
| H | 290 | 296 | 283 | 290 | 1159 |
| I | 550 | 650 | 650 | 650 | 2500 |
| J | 352 | 384 | 399 | 385 | 1520 |
| K | 175 | 300 | 300 | 125 | 900 |
| L | 750 | 900 | 899 | 890 | 3439 |
| M | 700 | 800 | 800 | 800 | 3100 |
| N | 602 | 693 | 695 | 700 | 2690 |
| O | 197 | 199 | 198 | 200 | 794 |
| P | 497 | 608 | 604 | 622 | 2331 |
| **Subtotal per insecticide** | 7553 | 8569 | 8559 | 8309 | Grand Total: 32990 |

### WHO tube bioassay results

Concentration-response curves for each insecticide and Kisumu colony are displayed in Figures 2-5. Descriptive analysis of all data, representing raw proportional values of mortality, including those datasets from which concentration-response curves could not be produced, are provided in the Supplementary Materials.

At four sites, an LC could not be estimated because testing was restricted to the control and the lowest and highest concentrations owing to resource constraints: sites F, H and O for permethrin, alpha-cypermethrin and DDT, and additionally site K for pirimiphos-methyl (site K tested two intermediate concentrations). For these sites, susceptibility was instead assessed from mortality at the discriminating concentration (DC), with mortality below 98% taken to indicate ’possible resistance’. Some variability was also seen between sites in mortality at the lowest concentration tested for each insecticide. Per-site LC estimates and DC mortality are given in Tables 3–6, with dose-response data in Figures 2–5.

**Table 3.** Predicted LC_50_ values for Kisumu exposed to a range of permethrin concentrations, across different testing sites. Sites with significantly different LC_50_ values are marked with different letters. Fold change is relative to the lowest LC_50_ estimate across the sites. Site K (*) two replicates were conducted out of the recommended temperature range during the exposure period: 1C out of range (too cool) at the exposure start and 0.8°C too high during the exposure end in another replicate). Site F (**), humidity was out of range (<60%) during holding and exposure periods in all replicates. Site O (***) did not run any controls

| Lab/Site | LC50 | Lower CI | Upper CI | Resistance/fold change | <100% mortality with DD |
| --- | --- | --- | --- | --- | --- |
| E | 0.0744<sup>a</sup> | 0.0662 | 0.0839 | 1 | N |
| C | 0.0903<sup>ab</sup> | 0.071 | 0.105 | 1.21 | N |
| J | 0.0929<sup>b</sup> | 0.0829 | 0.102 | 1.25 | N |
| L | 0.0937<sup>ab</sup> | 0.0509 | 0.101 | 1.26 | N |
| K* | 0.11<sup>c</sup> | 0.104 | 0.117 | 1.48 | - |
| P | 0.117<sup>c</sup> | 0.108 | 0.133 | 1.58 | N |
| G | 0.136<sup>d</sup> | 0.125 | 0.16 | 1.83 | N |
| I | 0.155<sup>de</sup> | 0.129 | 0.193 | 2.08 | N |
| B | 0.175<sup>def</sup> | 0.126 | 0.253 | 2.35 | Y |
| D | 0.178<sup>e</sup> | 0.158 | 0.198 | 2.4 | N |
| N | 0.213<sup>f</sup> | 0.198 | 0.23 | 2.87 | N |
| A | 0.225<sup>f</sup> | 0.195 | 0.248 | 3.02 | N |
| M | 0.29<sup>g</sup> | 0.251 | 0.338 | 3.9 | Y |
| F** | - | - | - | - | - |
| H | - | - | - | - | - |
| O*** | - | - | - | - | - |

**Table 4.** Predicted LC_50_ values for Kisumu exposed to a range of alpha-cypermethrin concentrations, across different testing sites. . Sites with significantly different LC_50_ values are marked with different letters. Fold change is relative to the lowest LC_50_ estimate across the sites. Site K (*) two replicates were conducted outside of the recommended temperature range during the exposure period: 1C out of range (too cool) at the exposure start and 0.8°C too high during the exposure end in another replicate). Site F (**), humidity was out of range (<60%) during holding and exposure periods in all replicates. Site O (***) did not run any controls

| Lab/Site | LC50 | Lower CI | Upper CI | Resistance/fold change | <100% mortality with DD |
| --- | --- | --- | --- | --- | --- |
| P | 5.15e-04<sup>a</sup> | 1.72e-04 | 1.04e-03 | 1 | N |
| L | 5.41e-04<sup>a</sup> | 1.85e-04 | 1.01e-03 | 1.05 | N |
| E | 6.57e-04<sup>a</sup> | 2.74e-04 | 1.09e-03 | 1.27 | N |
| N | 1.24e-03<sup>b</sup> | 8.78e-04 | 1.61e-03 | 2.41 | N |
| C | 1.49e-03<sup>b</sup> | 9.40e-04 | 2.27e-03 | 2.89 | N |
| D | 1.99e-03<sup>cd</sup> | 1.48e-03 | 2.65e-03 | 3.86 | N |
| K* | 2.17e-03<sup>cd</sup> | 1.48e-03 | 2.97e-03 | 4.21 | - |
| G | 2.19e-03<sup>c</sup> | 1.97e-03 | 2.50e-03 | 4.25 | N |
| A | 2.50e-03<sup>cd</sup> | 1.95e-03 | 3.22e-03 | 4.84 | N |
| J | 2.66e-03<sup>d</sup> | 2.32e-03 | 3.00e-03 | 5.16 | N |
| I | 4.97e-03<sup>e</sup> | 3.31e-03 | 8.03e-03 | 9.64 | N |
| B | 4.99e-03<sup>e</sup> | 2.83e-03 | 9.05e-03 | 9.68 | Y |
| M | 7.92e-03<sup>e</sup> | 6.69e-03 | 9.59e-03 | 15.37 | Y |
| O*** | - | - | - | - | N |
| H | - | - | - | - | Y |
| F** | - | - | - | - | N |

**Table 5.** Predicted LC_50_ values for Kisumu exposed to a range of DDT concentrations, across different testing sites. Sites with significantly different LC_50_ values are marked with different letters. Fold change is relative to the lowest LC_50_ estimate across the sites. Site K (*) two replicates were conducted out of the recommended temperature range during the exposure period: 1C out of range (too cool) at the exposure start and 0.8°C too high during the exposure end in another replicate). Site F (**), humidity was out of range (<60%) during holding and exposure periods in all replicates. Site O (***) did not run any controls

| Lab/Site | LC50 | Lower CI | Upper CI | Resistance/fold change | <100% mortality with DD |
| --- | --- | --- | --- | --- | --- |
| I | 0.14<sup>a</sup> | 0.0307 | 0.277 | 1 | Y |
| K* | 0.4<sup>b</sup> | 0.29 | 0.538 | 2.86 | N |
| G | 0.496<sup>b</sup> | 0.441 | 0.558 | 3.55 | N |
| L | 0.714<sup>c</sup> | 0.6 | 0.844 | 5.1 | Y |
| M | 0.746<sup>c</sup> | 0.619 | 0.905 | 5.34 | N |
| A | 0.786<sup>c</sup> | 0.604 | 1.01 | 5.62 | Y |
| N | 0.792<sup>c</sup> | 0.725 | 0.86 | 5.67 | N |
| D | 1.4<sup>d</sup> | 1.28 | 1.55 | 10.03 | Y |
| P | 1.44<sup>d</sup> | 1.31 | 1.56 | 10.28 | N |
| E | 1.61<sup>de</sup> | 1.39 | 1.92 | 11.51 | Y |
| C | 1.65<sup>de</sup> | 1.33 | 2.04 | 11.82 | Y |
| J | 1.81<sup>e</sup> | 1.57 | 2.11 | 12.96 | Y |
| B | 5.61<sup>f</sup> | 2.98 | 45.9 | 40.11 | Y |
| O*** | - | - | - | - | Y |
| H | - | - | - | - | N |
| F** | - | - | - | - | N |

**Table 6.** Predicted LC_50_ values for Kisumu exposed to a range of pirimiphos-methyl concentrations, across different testing sites. Sites with significantly different LC_50_ values are marked with different letters. Fold change is relative to the lowest LC_50_ estimate across the sites. Site K (*) two replicates were conducted out of the recommended temperature range during the exposure period: 1C out of range (too cool) at the exposure start and 0.8°C too high during the exposure end in another replicate). Site F (**), humidity was out of range (<60%) during holding and exposure periods in all replicates. Site O (***) did not run any controls

| Lab/Site | LC50 | Lower CI | Upper CI | Resistance/fold change | <100% mortality with DD |
| --- | --- | --- | --- | --- | --- |
| P | 6.96<sup>a</sup> | 4.12 | 9.64 | 1 | N |
| E | 7.42<sup>a</sup> | 4.35 | 11 | 1.07 | N |
| D | 8.55<sup>ab</sup> | 5.27 | 12.1 | 1.23 | N |
| C | 12.4<sup>b</sup> | 9.76 | 16 | 1.78 | N |
| B | 23<sup>c</sup> | 18.6 | 26.2 | 3.31 | N |
| A | 28.8<sup>d</sup> | 25.7 | 31.8 | 4.14 | N |
| L | 32.6<sup>e</sup> | 31.7 | 33.4 | 4.68 | N |
| M | 33<sup>e</sup> | 31.4 | 34.8 | 4.74 | N |
| N | 41.6<sup>f</sup> | 39.1 | 44.3 | 5.98 | N |
| K* | - | - | - | - | N |
| O*** | - | - | - | - | N |
| H | - | - | - | - | N |
| F** | - | - | - | - | N |
| G | - | - | - | - | N |
| I | - | - | - | - | N |
| J | - | - | - | - | N |

#### Permethrin

Estimated LC_50_ values for permethrin for Kisumu varied across different sites. The mean LC_50_ across sites was 0.15% (95% CI: 0.13%-0.17%), with the difference between the largest and smallest LC_50_ values equivalent to a fold change of 3.90 (Table 3). All excluded sites (F, H, O) showed 100% mortality at the DC, with the low-concentration variability most pronounced at site O. Possible resistance was indicated at sites B and M, both with <98% mortality at the DC (Table 3, Figure 2B).

#### Alpha-cypermethrin

Estimated LC_50_ values for alpha- cypermethrin for Kisumu varied across different sites. The mean LC_50_ across sites was 0.0026% (95% CI: 0.0019%-0.0037%), with the difference between the largest and smallest LC_50_ values equivalent to a fold change of 15.38 (Table 4). Of the excluded sites, F showed 100% mortality at the DC, while H and O averaged 97.8% and 99% respectively, the former indicative of possible resistance (Supplementary Materials). Possible resistance was also indicated at sites B and M, which showed marked survival at the DC (91% and 82% mortality respectively; Table 4, Figure 3).

#### DDT

Estimated LC_50_ values for DDT for Kisumu varied across different sites. The mean LC_50_ across sites was 1.35% (95% CI: 1.01%-4.62%), with the difference between the largest and smallest LC_50_ values equivalent to a fold change of 40.07 (Table 5). Of the excluded sites, F and H showed 100% mortality at the DC, while O averaged 97%, indicating possible resistance; low-concentration variability was again most marked at site O. Possible resistance was widespread for DDT, indicated at sites A–E, I, J, L and O, with DC mortality below 98% and falling as low as 24% at site B (Table 5, Figure 4).

#### Pirimiphos-methyl (PM)

Estimated LC_50_ values for PM for Kisumu varied across different sites. The mean LC_50_ across sites was 21.59 mg/m^2^ (95% CI: 18.89 mg/m^2^ -24.36 mg/m^2^), with the difference between the largest and smallest LC_50_ values equivalent to a fold change of 5.98 (Table 6). Mortality at the lowest concentration (10.65 mg/m²) varied at sites F, H and O, while all other sites reached 100%. All sites showed 100% mortality at the DC (170 mg/m²) and were therefore classified as susceptible (Table 6). Sites G, I and J reached 100% or near-100% mortality across all concentrations (Figure 5B).

### Correlation of resistance between insecticides

Despite all sites testing the same susceptible reference strain (Kisumu), predicted LC values varied substantially between sites for every insecticide, with fold-changes between the lowest and highest site estimates of up to 40.11× for DDT, 15.37× for alpha-cypermethrin, 3.9× for permethrin and 5.98× for pirimiphos-methyl (Tables 3–6). To test whether this between-site variation was consistent across insecticides we examined the rank correlation of site LC values between insecticides (Figure 6). Correlations between insecticides were generally weak and, given the small number of sites available per pair (n = 9–13), none reached statistical significance: the ranking of sites for one insecticide did not reliably predict their ranking for another (DDT vs alpha-cypermethrin ρ = −0.08, p = 0.792; DDT vs permethrin ρ = −0.27, p = 0.373; alpha-cypermethrin vs permethrin ρ = 0.48, p = 0.097). The strongest associations were a negative correlation between DDT and pirimiphos-methyl (ρ = −0.60, p = 0.097, n = 9) and positive correlations of permethrin with pirimiphos-methyl (ρ = 0.60, p = 0.097, n = 9) and alpha-cypermethrin with permethrin (ρ = 0.48, p = 0.097, n = 13), although none were significant. No insecticide pair showed the strong, consistent positive correlation that would indicate a stable site-level ranking of susceptibility. No correction for multiple comparisons was applied across the six pairwise tests, as the analysis is intended to be exploratory and hypothesis-generating rather than confirmatory; the p-values should therefore be interpreted as descriptive, particularly given the small number of sites per pair.

**Figure 6.**
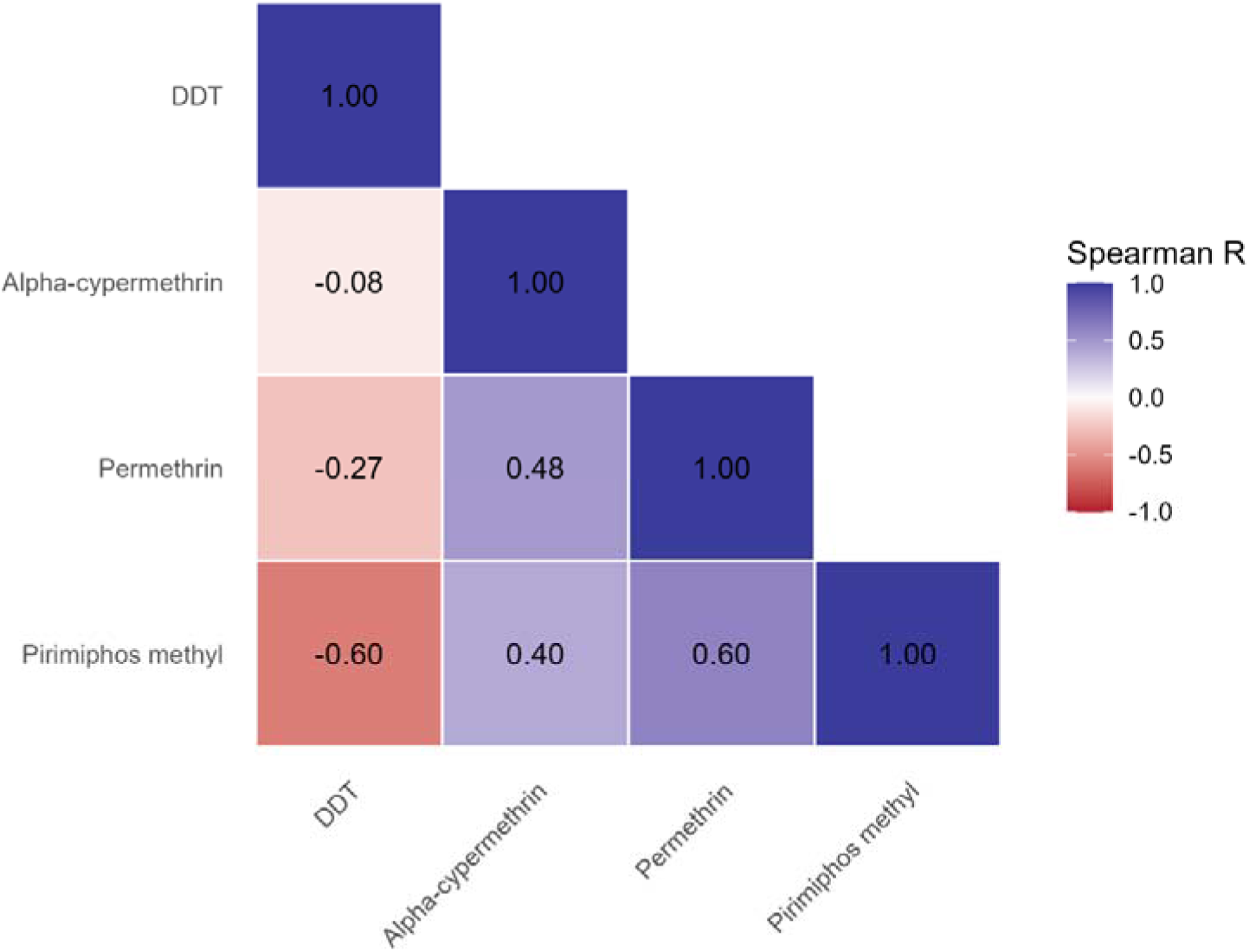
Spearman rank correlation (ρ) of site-level LC□□ estimates between insecticides for the Kisumu reference strain. Correlations were computed pairwise using all sites with an estimable LC□□ for both compounds (n = 9–13 per pair). Colour scale runs from −1 (red, perfect negative correlation) to +1 (blue, perfect positive correlation).

## Discussion

Well-characterised laboratory populations of insecticide-susceptible mosquitoes are widely used as reference comparators for evaluating insecticidal products and investigating insecticide resistance mechanisms. This practice assumes that a given reference strain produces consistent and comparable results across facilities and over time. The *Anopheles gambiae* Kisumu strain is one of the most commonly used reference populations for this purpose. This study characterised and compared the phenotypic susceptibility of Kisumu colonies held in sixteen laboratories worldwide to four insecticides, permethrin, alpha- cypermethrin, DDT and pirimiphos-methyl, and found that this assumption does not hold.

Substantial, insecticide-dependent differences in LC were observed between colonies, and some would be classified as resistant on the WHO discriminating-concentration criterion.

Such divergence is expected and mosquitoes, as a biological test system, are inherently variable in terms of bioassay results ^3,22^. Multi-centre studies have shown that colonies designated with the same strain name can differ substantially in their concentration– response profiles ^11,13^, likely reflecting the cumulative effects of long-term maintenance under different laboratory conditions. These include genetic drift, bottlenecks, inadvertent selection, and differences in rearing and husbandry practices, all of which can alter susceptibility even without intentional selection pressure. Accidental contamination with wild or resistant mosquitoes may also contribute to divergence. Consequently, strain identity alone is therefore not sufficient to guarantee equivalence between laboratories, and Kisumu held at different centres should be regarded as related but distinct laboratory populations. This highlights the value of profiling colonies phenotypically and genotypically ^6^ and presenting that characterisation alongside study results to improve the interpretation, reproducibility, and comparability of insecticide susceptibility studies ^23^.

Despite all sites testing the same susceptible reference strain (Kisumu), predicted LC values varied substantially between sites for every insecticide, with total site-to-site fold- changes ranging from 3.9× for permethrin to 40× for DDT (Tables 3–6). Mechan et al. ^3^ estimated a permethrin LC _50_ of 0.03% by probit analysis on Kisumu colony J (held at Liverpool Insect Testing Establishment, LITE) roughly three years earlier, against our estimate of 0.093% (95% CI 0.082–0.102%), a 3-fold difference. This illustrates the scale of variability surrounding LC estimates over time, which may not reflect a true change in susceptibility, as this colony is regularly profiled and characterised as susceptible ^6^. Most pairs of colonies differed significantly in their LC (95% credible intervals for the ratio excluding 1), indicating these are real differences rather than assay noise. This variation in WHO tube bioassay results between colonies and over time is consistent with the observation by Mechan et al.³ that roughly two-thirds of variation in mortality occurred between days within a single laboratory, suggesting that instability within a single colony over time adds to between-colony and between-laboratory differences as sources of variation.

The degree of divergence between colonies was strongly insecticide dependent but the majority of between-site differences were statistically supported: across the four insecticides, 76–86% of site pairs had LC -ratio credible intervals excluding 1 (panel C of Figures 2–5). Rather than forming a few discrete groups, sites generally lay along a graded susceptibility continuum in which adjacent sites overlapped but the extremes were clearly distinguishable. For permethrin, the spread was relatively contained (roughly 3.9-fold): the most susceptible sites (E, C, J, L) formed an overlapping cluster, while the least susceptible site (M) was distinguishable from every other site. Alpha-cypermethrin showed a wider spread (15-fold) with clearer separation at the extremes: the three most susceptible sites (P, L, E) formed a mutually indistinguishable group, distinct from a broad overlapping middle band and from the most resistant sites (I, B, M). Pirimiphos-methyl was similarly graded (6-fold), with 86% of site pairs distinguishable. DDT showed by far the greatest apparent divergence (40-fold), but this is likely an artefact rather than a biological signal and should be set aside when judging true between-colony variability. Risella oil, the carrier for DDT, is immiscible in acetone, making even coating of papers difficult and causing localised over- or under-dosing.

Consequently, DDT showed the greatest scatter of observed mortality around its fitted curves of the four compounds, exceeding 20% mean deviation at some sites. This is more variable than even the resistant strains characterised by Kont et al. ^21^ (≤9%), suggesting that the observed scatter was primarily methodological rather than biological origin. The 40-fold range is defined by the two least precisely estimated sites: the most resistant (B; 95% CI 2.98–45.9) and the most susceptible (I; 95% CI 0.03–0.28), both with credible intervals an order of magnitude wider than the other sites. Setting DDT aside, the remaining three compounds show moderate, graded variation rather than a few extreme outliers, with alpha-cypermethrin showing the clearest statistically supported divergence.

Placing this in context is possible because the same Bayesian concentration–response framework has now been applied to Kisumu in several multi-centre studies. In the WHO bottle bioassay study ^13^, Kisumu LC estimates for the pyrethroid transfluthrin varied roughly 4-fold across centres (comparable to our permethrin result) while other compounds showed between-centre ranges of around 9–10-fold (e.g. chlorfenapyr, clothianidin), with transfluthrin and flupyradifurone the most variable. Our alpha-cypermethrin result (15-fold) thus sits at or somewhat above the upper end of that study’s between-centre variation, while permethrin and pirimiphos-methyl are comparable or slightly below. A similar pattern emerged in establishing DCs for broflanilide and isocycloseram, where Kisumu LC estimates for isocycloseram spanned more than an order of magnitude across centres ^33^.

The same framework applied to laboratory strains used Kisumu as the susceptible reference against which resistant colonies were distinguished by one to two orders of magnitude; there, susceptible Kisumu showed low scatter around its fitted curve (∼3% mean deviation), comparable to our most-similar sites ^21^. Two caveats to these comparisons: those studies typically spanned only three centres per compound versus 9–13 here, so a larger raw range here partly reflects greater opportunity to capture extreme sites; and the comparisons span different assay formats (tube versus bottle), albeit analysed identically.

The finding that Kisumu colonies differ in measured susceptibility by several fold is consistent, but this variation is compound-specific rather than reflecting colonies that are uniformly more or less susceptible. Site rankings were largely inconsistent across insecticides: several colonies occupied opposite ends of the range depending on the compound (site I was most susceptible to DDT yet among the least to alpha-cypermethrin; site E was most susceptible to permethrin but among the least to DDT). The clearest reproducible signal was between the two pyrethroids, which share a target site and resistance mechanism: lab M was the least susceptible site for both, with narrow credible intervals indicating a genuine difference, and the most-susceptible sites overlapped (P, L and E low for both). Sites P and E ranked among the most susceptible across all three reliably estimated compounds, behaving as expected of a fully susceptible reference colony. That laboratory M was unremarkable for DDT and pirimiphos-methyl indicate that its elevated pyrethroid tolerance is a compound-specific effect rather than reflecting a generally less susceptible colony. The one partial exception to compound-specificity was site A, which ranked toward the less-susceptible end for all four compounds (ranks 12, 9, 6 and 6 of 13), making it the closest candidate for a broadly elevated tolerance that might warrant genotypic follow-up.

This site-level inconsistency is formalised by the rank-correlation analysis (Figure 6). If a shared mechanism drove the between-site variation we would expect positive correlations in site ranking (particularly between DDT and permethrin (which share kdr-mediated target-site resistance ^24^) and between the two pyrethroids, but this was not observed. The strongest single association, a negative DDT–pirimiphos-methyl correlation (ρ = −0.60), is mechanistically implausible and most likely reflects the lower reliability of the DDT assay.

Rather than a single shared driver such as a common resistance mechanism (e.g. kdr or P450-mediated metabolic resistance across these compounds ^24, 25^), the data point to compound-specific differences in tolerance, suggesting the colonies have diverged in response to specific local pressures rather than in overall susceptibility.

These differences carry broader implications, but the results also warrant caution for other reasons. Although all laboratories followed the same protocol and WHO tube SOP ^26^, slight differences in testing parameters and rearing conditions could not be fully disentangled from biological differences; environmental variation within acceptable ranges was not modelled, though a previous study found such fluctuations do not affect WHO tube results ^3^. Papers were treated by a single operator at one site ^27^, but without chemical confirmation of dose, and storage time before testing varied between laboratories.

The time between treating papers and using papers was also not consistent between laboratories, and the storage conditions before testing is a possible source of variability ^3^. Statistical limitations include the fact that an LC_50_ is inherently challenging to estimate ^28^, and that despite the sample size being specified in the protocol there was some variability in sample sizes actually tested between laboratories.

With those caveats, if colonies of a nominally single strain differ this much, several study types are affected: screening of new insecticides ^7^ risks false negatives if a colony carries cross-resistance; efficacy evaluations for prequalification, ^8,29,30^ may be underestimated; method-validation studies measuring inter-site variability ^32^ may be inflated; and transgenic work assuming a clean susceptible background ^33^, may be confounded. The clearest example is discriminating-concentration setting ^11,17,34^ which can be driven by the one or two least-susceptible sites: for broflanilide, LC estimates differed substantially between colonies labelled Kisumu, contributing to a several-fold increase in the proposed DC after multi-centre evaluation ^14,12^. Kisumu does not, therefore, represent a single uniform susceptibility phenotype.

These findings indicate that Kisumu and similar reference strains function as historical reference points rather than true biological standards. Continued reliance on strain name alone, without accounting for colony specific variation, risks introducing systematic inconsistency into, for example, discriminating concentration determination and resistance monitoring. Greater emphasis should therefore be placed on multi-centre validation and on recognising that susceptible reference strains represent laboratory-specific baselines rather than universal constants. Although Kisumu is among the most commonly-held susceptible strains of mosquito used in insecticide testing (9 of the 23 facilities participating in the WHO study to establish discriminating concentrations for new chemistries offered to test using Kisumu ^11^), other commonly used susceptible strains are Fang (*An. funestus*), and Rockefeller, Bora and New Orleans (*Ae. aegypti*). Whilst the focus of this study was the Kisumu strain, characterisation of additional significant susceptible reference strains should be considered.

A sample of mosquitoes was taken directly from each colony tested, and dead and alive mosquitoes were stored after the bioassay for future molecular analysis. Genetic analysis can therefore be performed to determine whether genetic differences between the colonies explain the differences in bioassay results. If not, then the remaining the variation in bioassay outcomes is due to differences in testing conditions, despite extensive efforts to standardise the resistance testing protocol across laboratories. If genetic differences exist, we can investigate their origin, using whole-genome sequencing to describe genetic variation across the colonies, identify any contamination with resistance haplotypes, and create a ‘family tree’ of Kisumu and determine whether systematic differences exist between different “branches” of the Kisumu colony’s history. If we are able to exclude genetic differences as the cause, we would have a measure of the inter-lab variability of the WHO tube bioassay which will be useful for interpretation of multi-site study data.

A molecular investigation of sub-colonies of the commonly used G3 susceptible strain of *An. gambiae* held in two laboratories discovered significant differentiation, with one strain containing more *An. gambiae* ancestry informative markers (AIMs) and the other strain more *An. coluzzii* AIMs ^35^. The strain was shown to have a mosaic genome, which illustrates how complicated it can be to attempt to mimic the responses of natural populations using a susceptible laboratory colony. A microsatellite analysis of fourteen laboratory colonies of *Aedes aegypti* ^36^ also found that colonies apparently of the same strain held in different laboratories were highly divergent, though less genetically variable than the field colonies to which they were compared. Differences in results were identified from virus competence studies using the same strain held at different laboratories, and the genetic composition of one strain was not compatible with what was thought to be the origin of one colony. These observations highlight the importance of maintaining careful colony records and characterising the mosquitoes used for research.

## Conclusion

This study compared the phenotypic insecticide susceptibility of sixteen colonies of the susceptible *Anopheles gambiae* “Kisumu” strain, held across laboratories worldwide, against four insecticides: permethrin, alpha-cypermethrin, DDT and pirimiphos-methyl. The results demonstrate that colonies of the *Anopheles gambiae* Kisumu strain maintained in different laboratories have diverged substantially in insecticide susceptibility, with some colonies exhibiting phenotypes consistent with resistance

These differences are potentially important because use of the Kisumu strain underpins a wide range of work in vector control research, including the screening and evaluation of new insecticides and products, the setting of discriminating concentrations, method validation, and the generation of transgenic lines. If colonies sharing a strain name diverge in susceptibility, the assumption that they provide a fixed, transferable baseline does not hold, and results which depend on that assumption may be biased or inconsistent between sites.

Possible explanations accounting for the observed divergence include genetic drift, bottlenecks, inadvertent selection, or contamination during long term colony maintenance, as well as differences in rearing conditions and unaccounted for variation in testing parameters. The weak and largely non-significant rank correlations between insecticides argue against a single shared resistance mechanism, pointing instead to compound-specific divergence between colonies. However, the relative contributions of genuine biological difference and residual methodological variation cannot be fully disentangled from the bioassay data alone, and genetic analysis of stored specimens is planned to further resolve this.

Overall, these findings indicate that reference strains like *Anopheles gambiae* Kisumu strains are best regarded as related but distinct laboratory-specific populations, functioning as historical reference points rather than universal biological standards. The difference between populations seen here underscores the importance of robust quality control measures, including careful colony record-keeping, rearing practices that minimise contamination, routine genotyping and phenotypic profiling, and full strain characterisation reported alongside study results, to support reliable interpretation and comparison of insecticide testing across laboratories.

## Declarations

### Data availability

This project contains the following underlying data: https://zenodo.org/records/20731457

Data are available under the terms of the Creative Commons Attribution 4.0 International license (CC-BY 4.0).

### Software availability

Source code available from: https://github.com/GioP93/What-is-Kisumu-Data

Source code is available under the terms of the GNU General Public License version 3 (GPL-3.0-only) (https://opensource.org/license/gpl-3-0).

### Author contributions

Gemma Francesca Harvey - Data Curation; Formal Analysis; Visualization; Writing – Original Draft Preparation; Writing – Review & Editing

Giorgio Praulins – Data Curation; Investigation; Project Administration; Formal Analysis; Visualization; Writing – Original Draft Preparation; Writing – Review & Editing

Frank Mechan – Formal Analysis; Visualization; Writing – Review & Editing

Alexandra Wright - Data Curation; Investigation; Methodology; Project Administration; Writing – Review & Editing

Rosemary Susan Lees – Conceptualization; Funding Acquisition; Methodology; Supervision; Writing – Original Draft Preparation; Writing – Review & Editing

## Acknowledgments

We would like to extend gratitude to Sarah Moore and Duncan Kobia Athinya for initial discussions which inspired this study. We thank Vestergaard Sarl for some initial discussions, and for contributing funding. We are very grateful to Agro & Life Solutions Research Laboratory, Sumitomo Chemical Co., Ltd. for providing data as one of the contributing laboratories.

